# Social plasticity in reproduction in female *Drosophila melanogaster*!

**DOI:** 10.64898/2025.12.20.695704

**Authors:** Simran Mann, Emily Rakosy, Sanduni Talagala, Tristan A.F. Long

## Abstract

The expression of behaviours in individuals are not always fixed and may vary according to the specific environmental conditions encountered by an individual. There is significant evidence that male fruit flies, *Drosophila melanogaster*, have the capacity to adjust their courtship behaviours depending on the social environment they encounter; however, the extent of plasticity in behavioural responses by females have received much less scrutiny. In this study, we set out to determine whether the presence and/or the types of conspecifics an individual female encounters influences her subsequent reproductive behaviour and patterns of egg production. In our assay, we housed medium-sized adult virgin females in experimental social treatments; either in isolation, or with other conspecific females (that were either of large- or small- body size) to observe how the presence and/or type of social interactions altered later reproductive decisions and outcomes. While in some instances female flies behaved differently depending on their housing treatment, these responses were not particularly consistent between flies obtained from different populations, nor with previously described patterns. Potential contributing factors for these results, and their implications for the future study of female reproductive plasticity are discussed.

## Introduction

The ability to respond plasticly to one’s environment is a widespread phenomenon which is thought to provide an individual with an adaptive benefit (Mery et al. 2010, Kasumovic and Brooks (2011), Snell-Rood 2013, Xue and Leibler. 2018, Fox et al. 2019, Yoon et al. 2023). Understanding how and why these responses are manifested provides insight into the relationship between the phenotype, the ecological context, and lifetime reproductive success. Considerable insights into this phenomenon have come from studies using fruit flies (*Drosophila melanogaster*), a species where behavioural, morphological and/or physiological traits are often shaped by their physical and/or social environment (Sheeba *et al*. 2002, Leadbeater 2009, Klepsatel *et al*. 2020, Chen and Sokolowski 2022, Schang *et al*. 2023, Shoot *et al*. 2024). In particular, many studies have focused on how male reproductive strategies change depending on their perception of their social environment, and what it may indicate about the current/future opportunities for mating and/or intra-sexual selection, and the conditions that their offspring may encounter (*e.g.* Friberg 2006, Bretman et al. 2010, Garbaczewska et al. 2013, Amos et al. 2025). However, much less in know about how (why, or even if) female reproduction is also influenced by the presence/absence/qualities of the same-sex conspecifics in their environment.

To begin to address this knowledge gap, Churchill *et al*. (2021) explored how female fruit flies altered their mating and ovipositing behaviours in response to their pre-mating social environment, as the choice of mate and the number and/or placement of her eggs can have important consequences for the success of their offspring (and ultimately their fitness) (*e.g.* Partridge 1980, Promislow et al. 1998, Bileter et al. 2012, Bettiet al. 2014, Talagala et al. 2024, Rakosy *et al*. 2025). In their study, recently eclosed virgin female *Canton-S* fruit flies were housed in isolation, or in groups for seven days, before to being used in mating assays. They observed that female flies that had been kept by themselves mated more quickly, mated for a longer period of time, and subsequently laid more eggs (in an ∼18-20h oviposition window) compared to those that had been previously housed with conspecifics. A similar response was also seen in a study by Fowler *et al*. (2022) where virgin *Dahomey* females were housed by themselves or with conspecifics (for varying durations of time), prior to mating assays, and it was also seen that solitary females mated more rapidly than socially-exposed females. Furthermore, more eggs were laid by the solitary females, and they mated for longer, than the socially-exposed females, but this difference between treatments was only seen when the social flies had been group-housed for over 48h. In both these studies the differences associated with social exposure were interpreted as a plastic response to the female’s different perceptions about their own future reproductive opportunities and/or the competitive developmental environment that their offspring might encounter in the future. Specifically, they hypothesized that group-housed females were more selective about immediately engaging in reproduction, if the presence of many conspecifics signalled that they would be more likely to encounter more (potentially high quality) males in the future and/or if they *did* choose to mate their offspring would be likely to encounter relatively higher levels of larval competition as a result of the simultaneous oviposition of the conspecifics in the same environment.

More recently, McConnell *et al*. (2024) investigated whether muti-generational selective manipulation of adult sex ratios (and thus the perceived availability of mates and/or the potential for offspring competition) affected the expression of social plasticity in reproduction. In their comparisons of social vs. group-housed females from experimentally evolved *Dhaomey*-derived populations that had been maintained in female-biased, male-biased or equal sex ratios, they saw the same pattern of mating latency plasticity as in Churchill *et al*. (2021), but no differences in mating duration in females from all selective treatment populations. Furthermore, a similar pattern of greater post-mating egg laying (over a ∼24h oviposition window) in the group-housed females compared to solitary females was also seen in all populations, irrespective of their selective history.

Together these (largely) consistent patterns are fascinating as they potentially allow us to better understand how females perceive and adjust their behaviours in responses to changes in their socio-sexual environment. However, much remains unknown about the factors that can might contribute to the expression of this phenomenon and the ways in which it may be manifested. As such, we set out to i) explore the extent to which the qualities (*i.e.* body size) of the conspecifics in the social groups influenced focal female, ii) measure the effects of offspring production over a longer post-mating time frame, iii) examine if plasticity is also manifested in egg qualities, and iv) determine if these patterns are seen in more populations of flies.

Our first objective involves experimentally manipulating the characteristics of the group-housed females’ social environment. If females are responding to the presence of conspecifics because their presence foretells future environments, the specific phenotypes of the conspecifics (and their cues) may be important. Previous studies, such as those by Cho *et al*. (2021) and Lin *et al*., (2021) have demonstrated that the specific characteristics of the conspecifics in a social group can influence the physiology of focal flies. We chose to vary the body size of the flies in the social group, as it is a phenotype that displays considerable variation (because of both genetic and environmental factors, *see* Hoffman *et al*. 1990, De Moed *et al*. 1997. Turner *et al*. 2011, Schang *et al*. 2023), and flies can perceive their own body size (Krause *et al*. 2019). Furthermore, as large-bodied females are more fecund, and more likely to be courted by males than small-bodied females (Long et al. 2009, Talagala *et al*. 2024, Freed *et al*. 2025), a focal female surrounded by larger conspecifics might anticipate a different potential future compared to a female whose social group is comprised of smaller individuals. Secondly, while these previous studies have focused on offspring production in the 18-24h period immediately after mating, it is unknown what might be manifested over a longer span of time. Thirdly, whether there are any changes to the quality of the eggs produced (measured as volume) has not been quantified. Finally, to better understand how universal this phenomenon is, it is necessary to conduct similar experiments in independent populations, with different life history schedules. Together our assays seek to better characterize the capacity and expression of socially-mediated plasticity in females, with the goal of understanding how social factors influence the expression of reproductive strategies in this important model species.

## Materials and Methods

### Population Protocols and Fly Maintenance

In our assays, we used focal fruit flies, *Drosophila melanogaster*, from two large, outbred *wild-type* laboratory populations: *Ives* (hereafter “IV”) and LHm, and social conspecific flies from two associated populations: IV-*bw* and LHm-*bw*. The IV population was initially established from a sample of 200 males and 200 females collected in Amherst, MA, USA in 1975 (Rose and Charlesworth 1981), while the LHm population originated from a sample of from 400 inseminated females obtained in an orchard near Modesto, CA, USA in 1991 (Rice *et al*. 2005). The IV-*bw* and LHm-*bw* populations were both created by introgressing the recessive allele, *bw*^1^ into the genomes of their *wild-type* counterparts via 10 rounds of back-crossing, to obtain populations that contain >99.9% of the IV or LHm genetic backgrounds, but where individuals express the *brown-eye* phenotype. We have subsequently performed additional rounds back-crossing in IV-*bw* and LHm-*bw* populations to minimize their divergence from their *wild-type* counterparts, with the last session occurring approximately 50 generations prior to the start of this experiment. We have maintained all four populations in our lab since 2011 in standard *Drosophila* vials (95mm H x 28.5mm OD) containing 10mL of banana/agar/killed-yeast food, dusted with live yeast (Rose 1984, Shoot *et al*. 2024) which are incubated at 25°C and 60% humidity on a 12L:12D hour diurnal light cycle. We have cultured these populations on non-overlapping 14-day generation schedules for hundreds of generations [for specific details on IV/IV-*bw*’s culturing protocols, see (Rose (1984); for LHm/LHm-*bw*’s see (Rice *et al*., 2005)].

### Collection of focal females and conspecific flies

From density-standardized vials of IV or LHm eggs, we collected focal females as virgins (within 8h of eclosion), under light CO_2_ anesthesia, and held them individually in test-tubes containing 1ml of standard media for 24-36h to allow them to complete their maturation and to confirm their unmated status (by the absence of larval production). We used a sieve sorter device (*see* Long *et al*. 2009) to quickly identify and isolate females of different size classes based on their ability (or inability) to pass through a column of sieves where the diameter of the electroformed holes decreased at each step by 5%. We anesthetized our collections of IV and LHm virgin females and (separately) sorted them by size and retained those individuals who were able to pass through the 1313µm diameter sieve, but were retained in the sieve with 1214µm diameter holes. These medium-sized focal females were then haphazardly placed into a new vial by themselves (“solitary” treatment) or onto one of two “social” treatment vials that contained five conspecifics from their corresponding, *brown-eyed* population that were either of a “large” or “small” phenotype. These social conspecifics females had been collected from density-standardized vials of IV-*bw* or LHm-*bw* eggs that had been established 2-3 days before the vials that yielded the *wild-type* focal females. Social females had also been collected as virgins and housed in sets of 10 for 24-36h before being categorized based on their sieve-size phenotype. We used the same size categories as in Talagala *et al*. (2024), where “large” individuals were defined as those that were too large to pass through the 1313µm sieve, while “small” females were defined as those that were small enough to pass through the 1167µm sieve. Once the 80 replicate vials for each of the solitary, “social large”, and “social small” treatments for each of the two populations had been assembled, they were returned to the incubator for 72h (a duration that Fowler *et al*. (2022) determined was sufficient for a social response to be expressed) prior to being used in the mating and offspring production assays.

### Mating and offspring production assays

We began the assay by lightly anesthetizing the female flies in their vials and transferring the focal female into a new vial containing fresh media and a similarly-aged *wild-type* virgin male (from the same population of origin as the focal female). We promptly mounted these vials horizontally in a quiet, well-lit room and observed for a period of 120 minutes. Trained observers who were nlind to the treatments, recorded the time of both the start and end of matings to the closest second. Vials were individually labelled, and observers were unaware of the specific treatment group that focal females belonged to.

Following the end of the observation period, the vials were left undisturbed for an additional 22h, at which time we removed the flies under light anesthesia, placed females into new vials (hereafter “Day 2” vials) and placed them and the “Day 1” vials into the incubator. After 24h, the females were removed from the “Day 2” vials and transferred into “Day 3” “mini-egg” chambers (*as in* Rakosy *et al*. 2025) which are constructed from polyethylene 20 mL sample vials (Kartell 733/4; 74.5mm H x 24.8mm OD) whose lid contained juice/agar media (Sullivan *et al*. 2000) and was used as a “dish” for ovipositing. We placed the mini-egg chambers and the “Day 2” vials in the incubator. Eighteen hours later we removed the focal females from the mini-egg chambers and counted the number of “Day 3” eggs that she had laid. These eggs were then gently aligned using a paintbrush dipped in embryo wash solution (Sullivan *et al*. 2000) into the center of the dish. We carefully placed a microscale, precise to 0.1mm (MT23, Electron Microscopy Sciences) next to the eggs, and photographed them using ScopeImage 9.0 image processing software (Microscopenet.com) and a 9.0MP trinocular-mounted camera. We used the image analysis program ImageJ (Schneider et al. 2012) with the ObjectJ plugin (Vischer & Nastase, 2022), to measure the eggs’ length (L) and width (W), which were used to calculate egg volume (V), using the formula for a prolate spheroid: V=1/6πW^2^L (*as in* Pitnick *et al*. 2003 and Pischedda *et al*. 2011).

The “Day 1” and the “Day 2” vials each remained in the incubator for 13 days at which time the number of adult flies present were counted and recorded.

### Statistical Analyses

All data analyses were conducted in the R statistical computing environment (version 4.0.3, R Core Team, 2020). For all analyses below we conducted tests of IV and LHm data separately and also created models where the independent factor either had three levels (“Solitary”, “Social Large” or “Social Small”) or two levels (“Solitary” or “Social”), where in the latter analyses, we pooled the data collected from our two social-exposure treatment groups.

In our analyses of mating behavioural phenotypes and their subsequent outcomes, we began by comparing the frequencies of successful copulations within the 90-minute observation window using Generalized Linear Models (GLMs), where the dependent variable was the observed copulation status, with a binomial error distribution. We determined whether the treatment means were significantly different from each other with likelihood-ratio chi-square tests using the *Anova* function in the *car* package (Fox and Weisberg 2011), which was followed, by calculating the estimated marginal means for each group, and (if necessary) post-hoc tests with Tukey method p-value adjustments using the *emmeans* function in the *emmeans* package (Lenth 2022) to determine the specific location of differences between groups.

For the analyses of mating latency (time from start of observation period until mating began) and mating duration (time between the start and end of mating), we constructed survival models using the *survreg* function in the *survival* package (Therneau 2022). For the mating latency data, flies that had not mated by the end of the observation period were assigned a value of 7200 seconds (120 minutes) and right censored. In each of our survival analyses, we initially created multiple alternative models in which we altered the specified error distribution (Weibull, logistic, gaussian, extreme, and exponential) and chose the model that yielded the lowest residual errors. Using the best fitting model, we then tested the significance of the treatment using the *Anova* function and conducted *post hoc* tests as described above with functions in the *emmeans* package.

We analysed the data on the number of offspring produced on Day 1, the number of offspring produced on Day 2, the combined number of offspring produced on Day 1 and Day 2, and the number of eggs produced on “Day 3” with GLMs, with either quassipoisson or poisson error distribution (depending on the specific distribution of the data).

As egg volume data were not normally distributed in all groups (determined using to Shaprio-Wilks tests) we used the non-parametric Kruskal-Wallis test (*kruskal.test*) to determine if there were any significant differences between treatments, and when needed, *post-hoc* testing was performed using the *kruskalmc* function in the *pgirmess* package (Giraudoux 2018).

## Results

When considering the observed mating rates of IV focal females (Figure 1a & 1c, Table 1a) we detected no significant differences when we compared the three social-exposure treatment groups (LRχ^2^=1.78, df=2, p=0.422), nor when the solitary female mating rates were compared to the rates of the combined socially-exposed focal females (LRχ^2^=0.44, df=1, p=0.505). Similarly, the mating latencies (Figure 1a & 1c, Table 2a) did not differ between groups whether the social-exposed females were treated separately (LRχ^2^=1.266, df=2, p=0.531) or combined (LRχ^2^=0.25, df=1, p=0.616). For the LHm females, there was a trend towards different rates of mating between the three groups (Figure 1b, Table 1b, LRχ^2^=0.08, df=2, p=0.531) that was driven by higher rates of mating in the focal females exposed to large conspecifics compared to the control/solitary females (*post-hoc comparison* p=0.09), while the small-conspecific exposed females exhibited an intermediate rate. When the two social groups were analysed together (Figure 1d, Table 1b), the combined rate of mating was also slightly higher than that of the control/solitary females (LRχ^2^=3.76, df=1, p=0.052). The mating latencies of the LHm focal females showed signs that social exposure treatment had an effect. As with the mating rates, solitary/control focal females were slower to mate than large-conspecific exposed females (LRχ^2^=5.40, df=2, p=0.07, *post-hoc comparison* p=0.06), and overall socially-exposed females were faster to mate than those in the solitary/control treatment (LRχ^2^=3.76, df=1, p=0.03). The durations of observed mating by focal females were not significantly associated with their social exposure treatment in either the IV (*Three group analysis*: LRχ^2^=0.64, df=2, p=0.42, Figure 2a; *Two group analysis*: LRχ^2^=0.99, df=1, p=0.61, Figure 2c, Table 3a) or the LHm (*Three group analysis*: LRχ^2^=1.62, df=2, p=0.44, Figure 2c; *Two group analysis*: LRχ^2^=1.45, df=1, p=0.23, Figure 2d, Table 3b) focal females.

**Figure 1.**
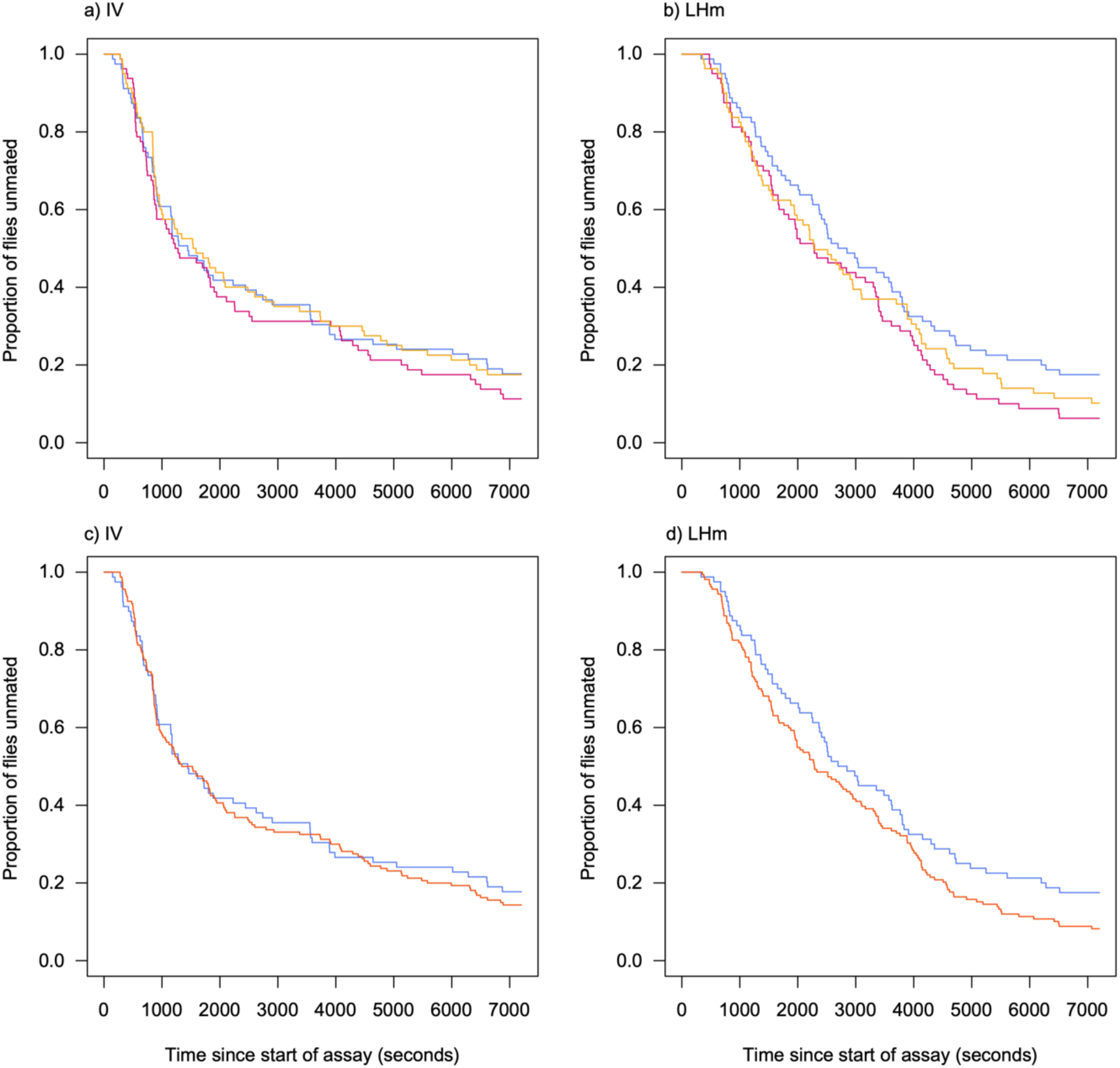
Survivorship curves depicting mating latencies (in seconds) of female fruit flies during the observation period. Figures in the left column (a & c) depict data obtained from flies from the IV population, while figures in the right column (b & d) depict data obtained from flies from the LHm population. In all figures data from flies in the “solitary/control” treatment are plotted with blue lines. In the top column figures (a & b), flies from the “large conspecific” treatment group are plotted with red lines, while flies from the “small conspecific” treatment group are plotted with yellow lines. In the bottom column figures (c & d) these two “social” treatments are pooled and plotted with orange lines.

**Table 1.**
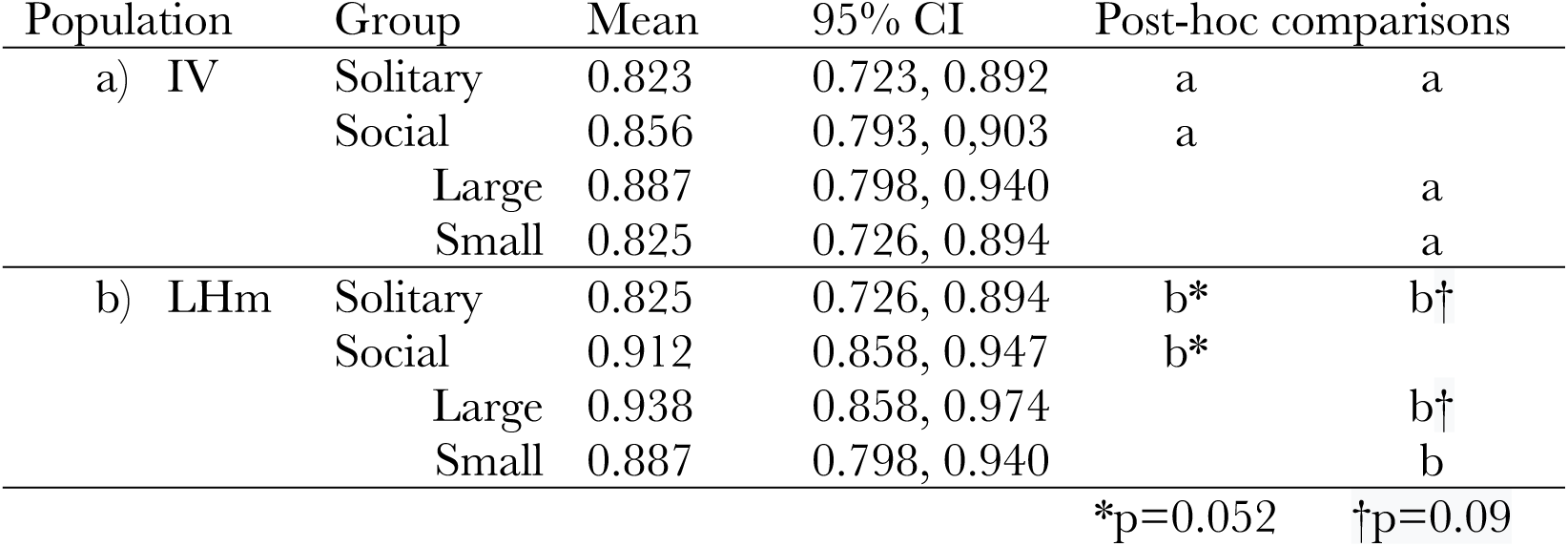
Mating Frequencies. Summary statistics of mating frequencies of flies from different social exposure treatments, and populations of origins that were observed to mate during the observation session. Groups with different letters are significantly different from each other according to post-hoc tests.

**Table 2.** Mating Latencies. Summary statistics of mating latencies (in seconds) of flies from different social exposure treatments, and populations of origins based on observations made during the observation session. Groups with different letters are significantly different from each other according to post-hoc tests.

| Population | Group | Mean | 95% CI | Post-hoc comparisons |  |
| --- | --- | --- | --- | --- | --- |
| a) IV | Solitary | 3354 | 2560, 4394 | a | a |
|  | Social | 3087 | 2561, 3722 | a |  |
|  | Large | 2807 | 2167, 3637 |  | a |
|  | Small | 3393 | 2597, 4434 |  | a |
| b) LHm | Solitary | 4219 | 3515, 5056 | b | b* |
|  | Social | 3329 | 2939, 3771 | c |  |
|  | Large | 3157 | 2942, 3750 |  | b* |
|  | Small | 3511 | 2942, 4189 |  | b |

\*p=0.06

**Figure 2.**
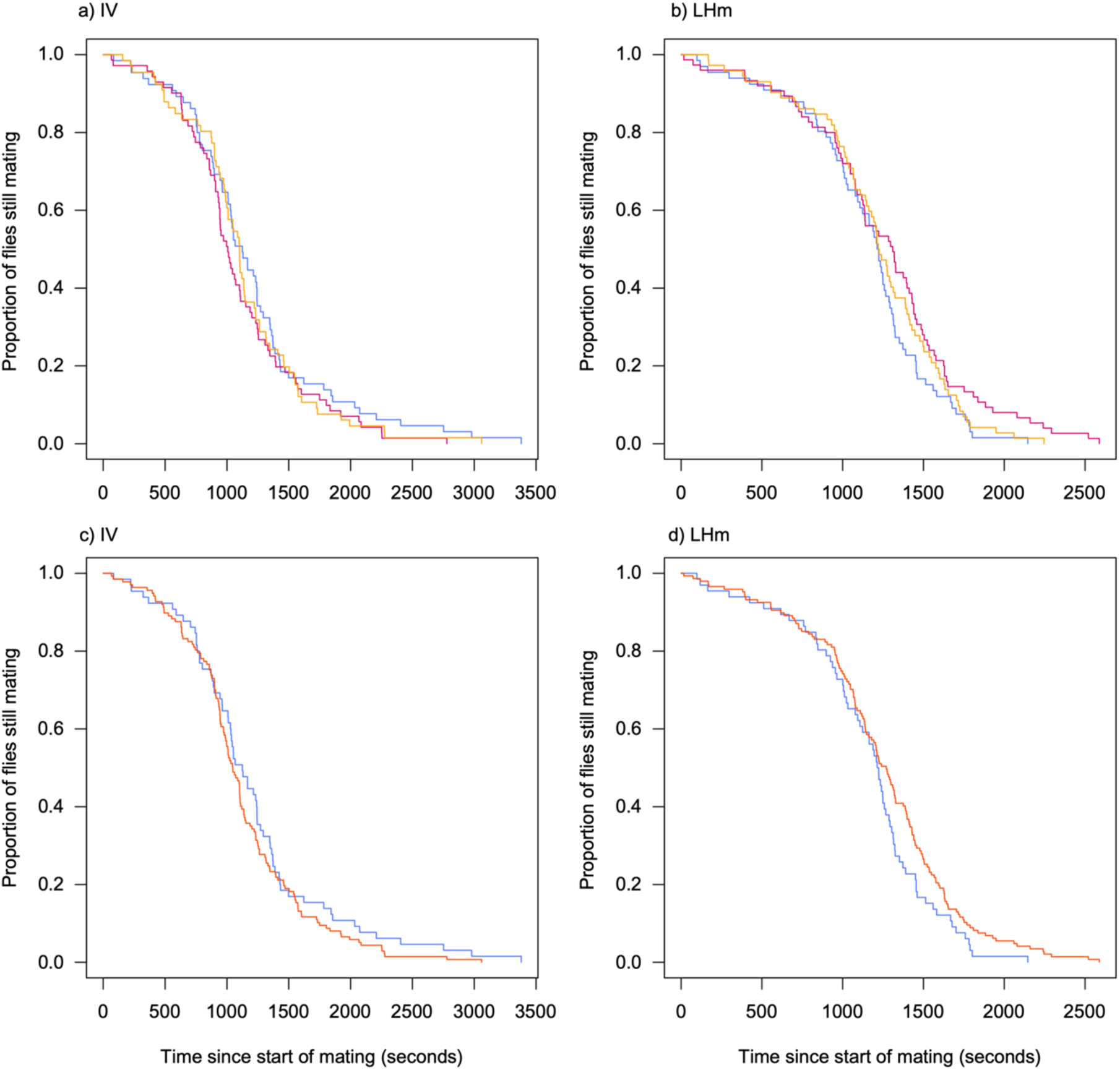
Survivorship curves depicting mating durations (in seconds) of female fruit flies seen copulating during the observation period. Figures in the left column (a & c) depict data obtained from flies from the IV population, while figures in the right column (b & d) depict data obtained from flies from the LHm population. In all figures data from flies in the “solitary/control” treatment are plotted with blue lines. In the top column figures (a & b), flies from the “large conspecific” treatment group are plotted with red lines, while flies from the “small conspecific” treatment group are plotted with yellow lines. In the bottom column figures (c & d) these two “social” treatments are pooled and plotted with orange lines.

**Table 3.** Mating Durations. Summary statistics of mating durations (in seconds) of flies from different social exposure treatments, and populations of origins that were observed to mate during the observation session. Groups with different letters are significantly different from each other according to post-hoc tests.

| Population | Group | Mean | 95% CI | Post-hoc comparisons |  |
| --- | --- | --- | --- | --- | --- |
| a) IV | Solitary | 1140 | 1021, 1258 | a | a |
|  | Social | 1082 | 1002, 1162 | a |  |
|  | Large | 1058 | 947, 1170 |  | a |
|  | Small | 1106 | 991, 1220 |  | a |
| b) LHm | Solitary | 1176 | 1075, 1277 | b | b |
|  | Social | 1251 | 1181, 1321 | b |  |
|  | Large | 1266 | 1166, 1367 |  | b |
|  | Small | 1236 | 1139, 1334 |  | b |

In the days that followed the initial male exposure of focal females we measured offspring and egg production. In the IV focal females, socially-exposed females laid more eggs laid than those in the solitary/control treatment on the first day of the assay (Figure 3e, LRχ^2^=9.00, df=1, p=0.003), which was largely driven by small-exposed females producing more offspring than, solitary/control focal females (Figure 3a, LRχ^2^=10.72, df=2, p=0.005, *post-hoc comparison* p=0.004, Table 4a). On the subsequent day of the experiment, there were no statistically significant differences in offspring production between groups, either in the three group analysis (LRχ^2^=2.40, df=2, p=0.30, Figure 3b, Table 4b) nor in the two group comparison (LRχ^2^=2.38, df=1, p=0.123, Figure 3f, Table 4b). Cumulatively over the first two days of the assay the social-exposed females produced ∼28% more offspring than the control/solitary females (LRχ^2^=7.91, df=1, p=0.005, Figure 3g, Table 4c), which was driven by the higher offspring production of small-conspecific-exposed females relative to the control/solitary focal females (LRχ^2^=8.92, df=2, p=0.01, *post-hoc comparison* p= 0.0095, Figure 3c, Table 4c), with large-exposed focal females producing an intermediate number of offspring over this period of time. For LHm focal females, we also detected a significant difference between the number of offspring produced during the first day of the assay between the socially-exposed and solitary/control group (LRχ^2^=4.50, df=1, p=0.03), but unlike in the IV focal females, the latter group produced more offspring than the former (Figure 4e, Table 5a). When offspring production was compared between the three groups we also detected significant differences between treatments (LRχ^2^=13.55, df=2, p=0.001), which was due to the small-exposed focal females producing fewer offspring than the solitary/control group (*post-hoc comparison* p=0.002) *and* fewer offspring than the large-exposed females (*post-hoc comparison* p=0.008). On the second day of the experiment, however there were no statistically significant differences in offspring production between groups for either the three-group analysis (LRχ^2^=0.4.27, df=2, p=0.11, Figure 4b, Table 5b) nor in the two-group comparison (LRχ^2^=0.14, df=1, p=0.71, Figure 4f, Table 5b). Cumulatively over the first two days of the assay there were no statistically significant differences in offspring production between groups for either the three-group analysis (LRχ^2^=0.4.27, df=2, p=0.11, Figure 4c, Table 5c) nor in the two-group comparison (LRχ^2^=1.87, df=1, p=0.17, Figure 4g, Table 5c).

**Figure 3.**
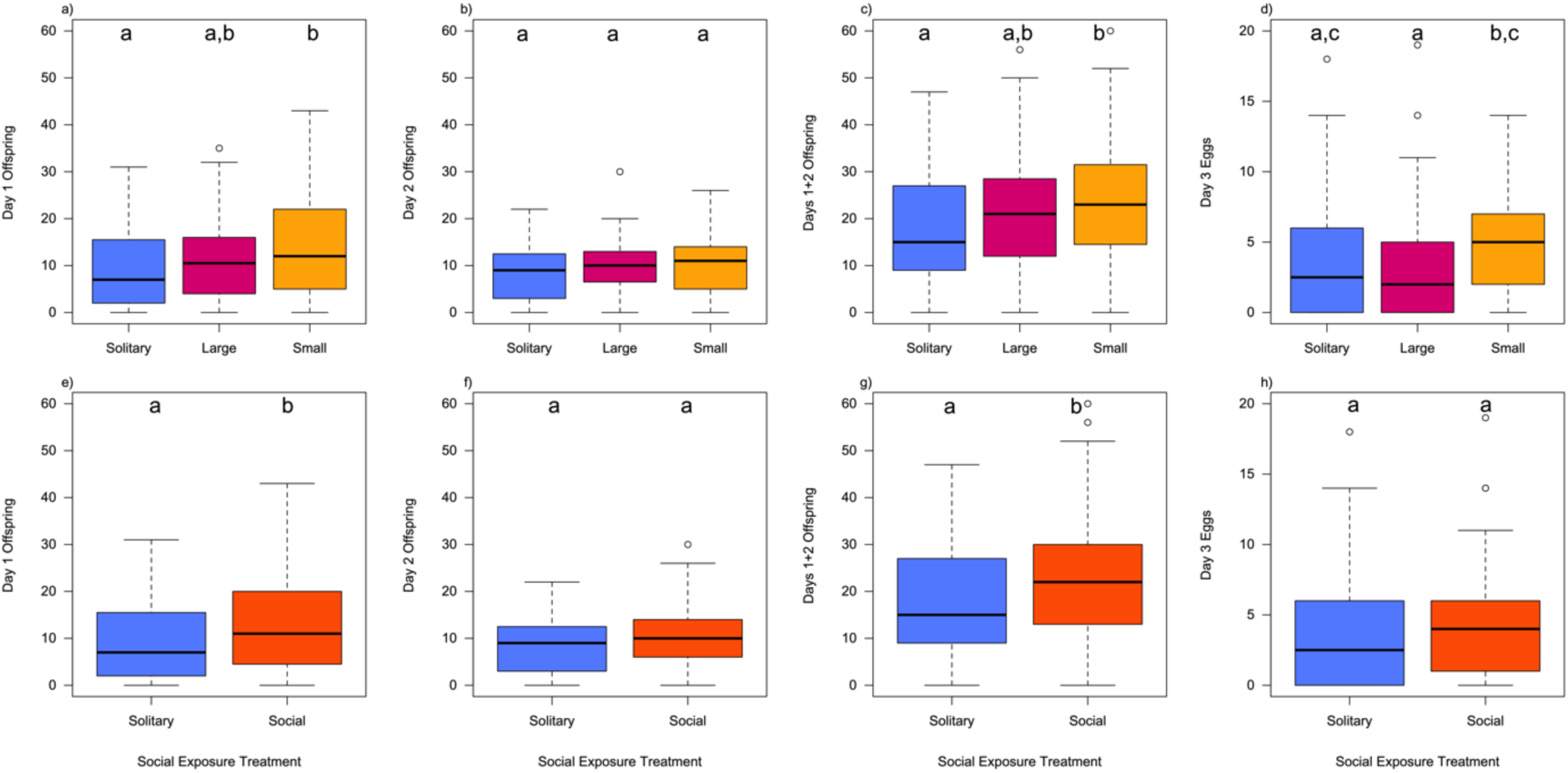
Boxplots depicting offspring and egg production of female flies from the IV population at different points in the assay. The boxes indicate the middle 50% of data (inter-quartile range, IQR), with the black horizontal line representing the median. Data points that are > ±1.5*IQR are outliers and indicated by circles. Whiskers extend to largest/smallest values that are not outliers. Figures in the first column (a & e) depict offspring production in the first 24h of the assay, those in the second column (b & f) depict offspring production in the second 24h of the assay, those in the third column (c & g) depict cumulative offspring production over 48h of the assay, and those in the fourth column (d & h) depict egg production in the final 18h of the assay. In all figures data from flies in the “solitary/control” treatment are depicted with blue. In the top column figures (a, b, c & d), flies from the “large conspecific” treatment group are depicted with red, while flies from the “small conspecific” treatment group are depicted as yellow. In the bottom column figures (e, f, g & h) these two “social” treatments are pooled and plotted in orange. Significant differences between groups indicated by post hoc lettering above each boxplot.

**Figure 4.**
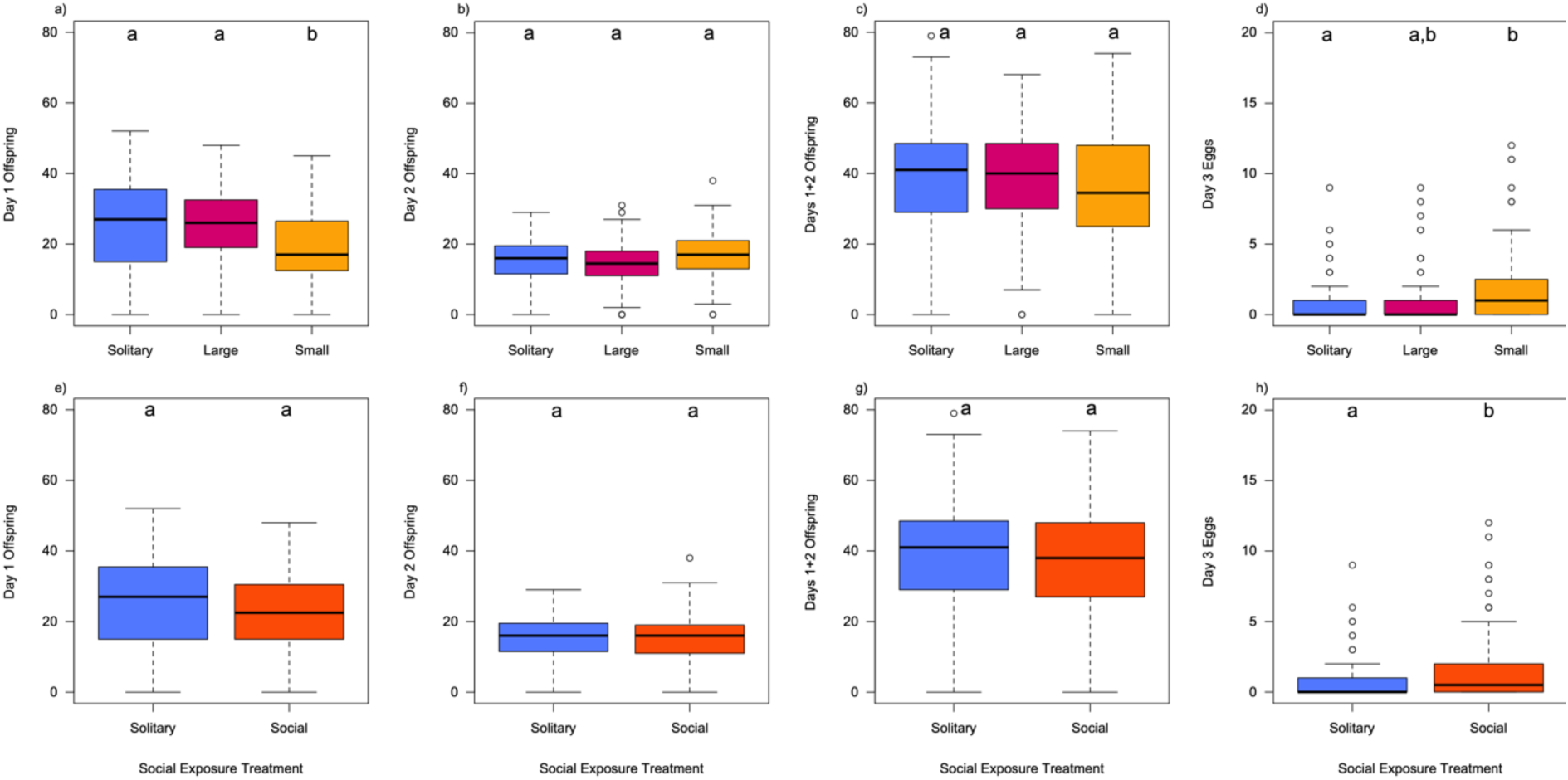
Boxplots depicting offspring and egg production of female flies from the LHm population at different points in the assay. The boxes indicate the middle 50% of data (inter-quartile range, IQR), with the black horizontal line representing the median. Data points that are > ±1.5*IQR are outliers and indicated by circles. Whiskers extend to largest/smallest values that are not outliers. Figures in the first column (a & e) depict offspring production in the first 24h of the assay, those in the second column (b & f) depict offspring production in the second 24h of the assay, those in the third column (c & g) depict cumulative offspring production over 48h of the assay, and those in the fourth column (d & h) depict egg production in the final 18h of the assay. In all figures data from flies in the “solitary/control” treatment are depicted with blue. In the top column figures (a, b, c & d), flies from the “large conspecific” treatment group are depicted with red, while flies from the “small conspecific” treatment group are depicted as yellow. In the bottom column figures (e, f, g & h) these two “social” treatments are pooled and plotted in orange. Significant differences between groups indicated by post hoc lettering above each boxplot (if applicable).

**Table 4.** IV Reproductive Output. Summary statistics of offspring production over the first 2 days of the assay, and egg production on the 3^rd^ day of the assay by flies originating from the IV population that experienced different social exposure treatments. Groups with different letters are significantly different from each other according to post-hoc tests.

| Variable | Group | Mean | 95% CI | Post-hoc comparisons |  |
| --- | --- | --- | --- | --- | --- |
| a) Day 1 Offspring | Solitary | 8.99 | 7.33, 11.0 | a | a |
|  | Social | 12.80 | 11.34, 14.4 | b |  |
|  | *Large* | 11.79 | 9.87, 14.1 |  | a,b |
|  | *Small* | 13.81 | 11.72, 16.3 |  | b |
| b) Day 2 Offspring | Solitary | 8.49 | 7.34, 9.81 | a | a |
|  | Social | 9.72 | 8.83, 10.7 | a |  |
|  | *Large* | 9.64 | 8.41, 11.04 |  | a |
|  | *Small* | 9.80 | 8.56, 11.22 |  | a |
| c) Days 1+2 Offspring | Solitary | 17.5 | 15.0, 20.4 | a | a |
|  | Social | 22.5 | 20.5, 24.8 | b |  |
|  | *Large* | 21.4 | 18.7, 24.6 |  | a,b |
|  | *Small* | 23.6 | 20.7, 26.9 |  | b |
| d) Day3 Eggs | Solitary | 3.52 | 2.84, 4.39 | a | a,c* |
|  | Social | 4.06 | 3.52, 4.69 | a |  |
|  | *Large* | 3.23 | 2.57, 4.05 |  | a |
|  | *Small* | 4.91 | 4.08, 5.91 |  | b,c* |

\*p=0.06

**Table 5.** LHm Reproductive Output. Summary statistics of offspring production over the first 2 days of the assay, and egg production on the 3^rd^ day of the assay by flies originating from the LHm population that experienced different social exposure treatments. Groups with different letters are significantly different from each other according to post-hoc tests.

| Variable | Group | Mean | 95% CI | Post-hoc comparisons |  |
| --- | --- | --- | --- | --- | --- |
| a) Day 1 Offspring | Solitary | 25.5 | 22.9, 28.4 | a | a |
|  | Social | 22.0 | 20.3, 23.9 | b |  |
|  | *Large* | 24.7 | 22.2, 27.5 |  | a |
|  | *Small* | 19.3 | 17.1, 21.8 |  | b |
| b) Day 2 Offspring | Solitary | 14.8 | 13.4, 16.5 | a | a |
|  | Social | 15.2 | 14.2, 16.3 | a |  |
|  | *Large* | 14.1 | 12.7, 15.6 |  | a |
|  | *Small* | 16.3 | 14.8, 18.0 |  | a |
| c) Days 1+2 Offspring | Solitary | 40.4 | 36.8, 44.3 | a | a |
|  | Social | 37.2 | 34.8, 39.9 | a |  |
|  | *Large* | 38.8 | 35.3, 42.7 |  | a |
|  | *Small* | 35.6 | 32.3, 39.3 |  | a |
| d) Day3 Eggs | Solitary | 0.80 | 0.50, 1.29 | a | a |
|  | Social | 1.49 | 1.17, 1.91 | b |  |
|  | *Large* | 1.16 | 0.79, 1.71 |  | a,b |
|  | *Small* | 1.82 | 1.34, 2.49 |  | b |

We counted and measured eggs produced by focal females on the third day of the assay. For IV focal females, while there were no overall statistically significant differences in egg numbers produced by the social and the solitary/control groups (LRχ^2^=1.17, df=1, p=0.28; Figure 3h, Table 4d), there was significant heterogeny across the three social exposure treatment groups (LRχ^2^=9.21, df=2, p=0.01, Figure 3d, Table 4d), with the small-conspecific exposed focal females laying significantly more eggs than the large-exposed focal females (*post-hoc comparison* p=0.01), and marginally more eggs than the solitary/control females (*post-hoc comparison* p=0.06). The measured volume of eggs produced on the third day of the assay by IV focal females was not significantly different when the grouped socially-exposed females were compared to the solitary/control females (Kruskall-Wallis χ^2^=1.02, p=0.31, Figure 5c), but significant heterogeny in egg volume was detected in the three-group analysis (Kruskall-Wallis χ^2^=0.25, p=0.0007), wherein small-exposed focal females laid larger eggs than either the large-exposed or the solitary/control females (Figure 5a, Table 6a). For LHm focal females, socially-exposed focal females laid more eggs than the solitary/control females (LRχ^2^=5.87, df=1, p=0.02, Figure 4h, Table 5d), which was driven by differences between the three experimental groups (LRχ2=9.33, df=2, p=0.009), wherein small-exposed females laid significantly more eggs than the control/solitary females (*post-hoc comparison* p=0.06), while the females exposed to large conspecifics produced intermediate numbers of eggs (Figure 4d, Table 5d). The measured volume of eggs produced on the third day of the assay by LHm focal females were not significantly different when the grouped socially-exposed females were compared to the solitary/control females (Kruskall-Wallis χ^2^=0.25, p=0.62, Figure 5d, Table 6b), nor when the three groups were compared with each other (Kruskall-Wallis χ^2^=0.28, p=0.87, Figure 5c, Table 6b).

**Figure 5.**
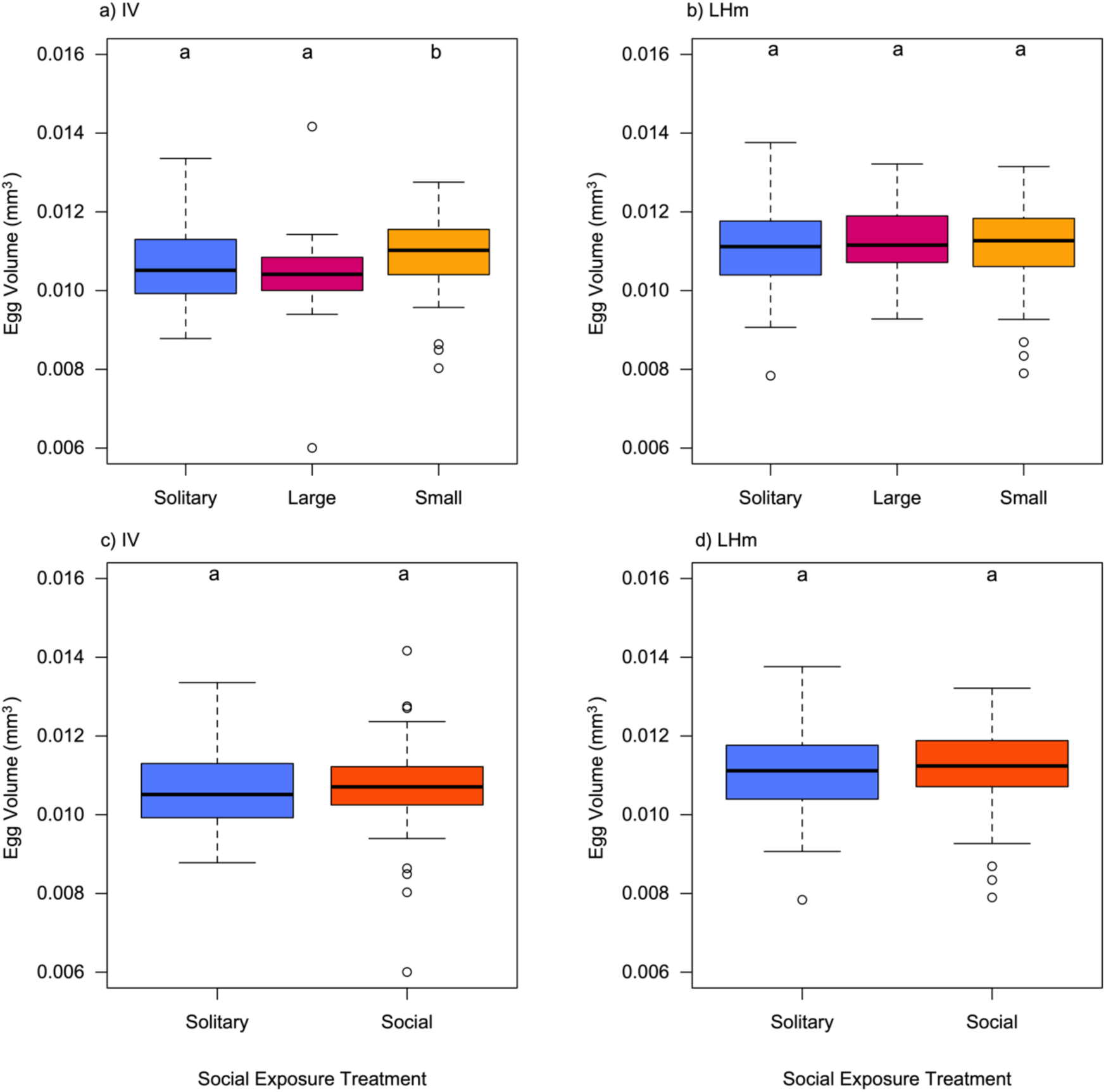
Boxplots depicting mean volumes (in mm^3^) of eggs laid by female flies on the third day of the assay. The boxes indicate the middle 50% of data (inter-quartile range, IQR), with the black horizontal line representing the median. Data points that are > ±1.5*IQR are outliers and indicated by circles. Whiskers extend to largest/smallest values that are not outliers. Figures in the left column (a & c) depict data obtained from flies from the IV population, while figures in the right column (b & d) depict data obtained from flies from the LHm population. In all figures data from flies in the “solitary/control” treatment are depicted with blue. In the top column figures (a & b), flies from the “large conspecific” treatment group are depicted with red, while flies from the “small conspecific” treatment group are depicted as yellow. In the bottom column figures (c & d) these two “social” treatments are pooled and plotted in orange. Significant differences between groups indicated by post hoc lettering above each boxplot (if applicable).

**Table 6.** Egg Volume. Summary statistics of average volume of eggs (in mm^3^) laid by flies from different social exposure treatments, and populations of origins. Groups with different letters are significantly different from each other according to post-hoc tests.

| Population | Group | Pseudo-<br>median | 95% CI | Post-hoc<br>comparisons |  |
| --- | --- | --- | --- | --- | --- |
| a) IV | Solitary | 0.01058 | 0.01029, 0.01086 | a | a |
|  | Social | 0.01076 | 0.01061, 0.01089 | a |  |
|  | *Large* | 0.01042 | 0.01023, 0.01060 |  | a |
|  | *Small* | 0.01099 | 0.01078, 0.01118 |  | b |
| b) LHm | Solitary | 0.01111 | 0.01060, 0.01160 | c | c |
|  | Social | 0.01123 | 0.01100, 0.01149 | c |  |
|  | *Large* | 0.01128 | 0.01090, 0.01167 |  | c |
|  | *Small* | 0.01122 | 0.01082, 0.01152 |  | c |

## Discussion

The presence and/or type of social cues encountered by an individual can result in the expression of plastic responses that lead to changes in behavioural, morphological and/or physiological phenotypes in order to better meet their specific environmental conditions, and ultimately increase their lifetime reproductive success (i.e., fitness) (Dore et al., 2018; Price et al., 2003). While it has been observed that changes in social environment can indirectly influence decision-making behaviors in *Drosophila melanogaster* during mating and offspring production, the study of behavioural plasticity has been much more extensively researched in males than in females (Churchill et al., 2021). Our study set out to examine i) if females respond with plastic mating behaviors to the presence of conspecifics and ii) if their response depended on the size of conspecifics they interacted with, and iii), whether the social cues generated by varying conspecifics influence a focal female’s egg production and quality. While we observed some cases where focal female flies responded differently to social isolation versus being co-housed with large- or small-bodied conspecifics, in many cases no statistically significant differences were observed, and the cases of differential patterns of response were frequently not consistent between our two test populations (or with previously published studies). Below we explore these finding and speculate on their implication for future research on this interesting field of inquiry.

The decision whether to mate or not can potentially be influenced by any number of (not necessarily independent) factors (Jennions and Petrie 1997, Billeter *et al*. 2012, Candolin 2019). If the perception of current and/or future reproductive opportunities is important to fruit flies, one would predict that cues obtained from conspecifics (or the lack thereof in their absence) might affect how females adjust their mating behaviour. In Churchill *et al*. (2021) the longer mating latencies and shorter copulation durations of their group-housed females, compared to their solitary flies, was interpreted as the former group exhibiting greater choosiness resulting from the inference that there would be more mating opportunities in the future (while also conceding that it was equally plausible that encountering only female conspecifics could also be interpreted by the flies as a scarcity of potential mating opportunities, in which case the solitary flies might end up exhibiting greater choosiness). In our assays we either saw no significant differences between the social and solitary conditions (in the case of the IV focal females), or higher/faster rates of mating (overall) in the socially-housed LHm focal females compared to the solitary/control focal flies. In these latter cases the differences between the solitary and social group means were largely driven by the responses of focal females housed with large-bodied conspecifics, with the focal females co-housed with small-bodied females behaving in a manner that was not statistically different from the control/solitary focal flies. We had initially hypothesized that the phenotype of the co-housed conspecifics might cause focal flies to behave differently as female body-size is correlated with both potential fecundity and reproductive interest from males (Long *et al*. 2009, Talagala *et al*. 2024) and focal females housed with large-bodied females might infer that they may be courted less than their conspecifics, and furthermore their offspring may encounter greater levels of competition during larval development. As fruit flies are capable of perceiving their own body size (Krause *et al*. 2019), and male fruit flies can distinguish between large- and small-bodied females, (Long *et al*. 2009, Lev and Pischedda 2023), we presumed that female flies would be able to perceive the phenotypic size of conspecifics, be it visually or indirectly via the amount of cuticle hydrocarbons (CHCs) that they expressed (*see* Luo *et al*. 2019). While the patterns of mating we observed were consistent with an inference of decreased choosiness following co-housing with large-bodied females, it could also potentially result from changes in the phenotype of the females as a result of their co-habitation with large-bodied females that made them more attractive to the mates. The transfer of CHCs from one female to another can change how they are perceived by males (*e.g*. Friberg 2006, Thomas 2011). It is conceivable that the increased mating rates are a result of the males being more motivated to court and mate the focal flies that had been co-housed with large-bodied females as they smelled like their conspecifics (a response that did not however translate into differences in copulation durations), and is a topic that is worth further investigation. In any event, while we did see differences in the LHm focal flies, the lack of a response in the IV focal females, coupled with no differences between any groups for mating duration indicates that reproductive plasticity in female *D. melanogaster* is more complicated and heterogenous than has been previously realized.

The second objective of this study was to examine the reproductive consequence (if any) of pre-mating social housing conditions. In previous studies by Churchill *et al*. (2021), Fowler *et al*. (2022) and McConnell *et al*. (2024), socially isolated females all laid more eggs and/or produced more progeny than their co-housed counterparts in the 1^st^ 24h following mating. While we did observe a similar difference when examining offspring produced in the 1^st^ 24h by the LHm focal females, in the case of the IV focal females the opposite pattern of offspring production was observed, wherein more offspring were produced by co-housed flies than those that had been kept in isolation. Interestingly in both cases the focal females housed with small-bodies conspecifics showed a *greater* plastic response than those that had been housed with large-bodied conspecifics. In Churchill *et al*. (2021), it was speculated that a reduction in egg production (in their case for females in the socially-exposed treatment) might be a response to avoid perceived higher rates of larval competition that might be encountered by their offspring in the future. Since the medium-bodied focal flies in our social-small treatment vials were surrounded by females of relatively lesser capacity for egg-production, it is conceivable that these focal females did not infer a low future competition relative to those focal females in the large-bodied treatment vials (or the females in the control/solitary vials). Our study also looked at egg/offspring production beyond the first 24h post-mating, which was a topic that had not previously been explored. In neither our IV, nor our LHm focal females, did we observe any significant differences in offspring production in the “Day 2” vials between any of the treatment groups. While this might suggest that the plastic effects of social exposure are thus manifested over a brief time frame, in the “Day 3” egg counts we did observe that for the IV focal females, that large-bodied exposed focal females produced fewer eggs than small-bodied exposed females (with the control/solitary focal females producing an intermediate number of eggs, and similarly in the LHm focal females the greatest number of eggs were produced by flies that had been co-housed with small-bodied conspecifics. Interestingly it was the females in the small-bodied social treatment that also produced the most voluminous eggs (at least in the case of the IV-derived focal females), which indicates there were no apparent trade-offs of quantity vs quality of eggs for the flies that produced eggs on the 3^rd^ day of the assay. While egg volume can be influenced by the genotype of the male with whom she mates (Pischedda *et al*. 2011), and is correlated with differences in female body size (TAFL and R. Hotwani, *unpublished results*), since single males were assigned haphazardly to females of standardized-size, it is unlikely that these factors may have contributed to our observed results. Together, these results suggest that any plastic responses to pre-mating social environmental conditions may be manifested on a longer time-frame than has been previously appreciated, and also in ways that were unexpected.

Given the notable differences in our results to those of previous studies by Churchill *et al*. (2021), Fowler *et al*. (2022) and McConnell *et al*. (2024), it is worth considering why such differences may have arisen, as well as what this may mean for future studies. While IV, LHm, *Dahomey* and *Canton-S* all have different origins and have been maintained in the lab on different diets, perhaps the most notable difference is that IV and LHm have always been maintained in vials of non-overlapping generations, while *Dahomey* and *Canton-S* are frequently cultured in population cages, with overlapping generations. One might speculate that in our study’s two populations, where at the start of each generation the age of each cohort is standardized in time and in numbers that the resulting synchronization of development and eclosion means that for hundreds upon hundreds of generations that adult flies rarely find themselves either alone (for very long) or surrounded by conspecifics of consistently larger- or smaller-body size. In contrast, in cage populations, where different cohorts of flies laid at different points in time may eclose asynchronously, and the population size may fluctuate temporally, there may be more instances when adult flies will find themselves in an environment that where conspecifics are either rare, or common. Thus, non-overlapping generations could conceivably lead to the loss of any ancestral capacity for plasticity in these lab-adapted populations. While this may be plausible, it should be noted that McConnell *et al*. (2024) used experimentally evolved lines created by Rostant *et al*. 2020 in which flies that originated from a *Dahomey* population cage were subsequently cultured for over 100 generations with non-overlapping generations and still exhibited a consistent plastic response to social manipulation.

While is it impossible to tell for certain why our results were different from those reported in other studies (and also between flies our two populations), our results do suggest that the expression of reproductive plasticity in female *Drosophila melanogaster*, a recently described phenomenon, is surprisingly variable and somewhat unpredictable. Future studies designed to understand what factors that contribute to its expression (or lack thereof), will be important to determine how this plasticity may contribute to variation in individual fitness and ultimately the evolution of species.

## Acknowledgements

We would like to thank Lukas Ghiglione, and Harleen Taneja of the #DrosLife Lab for their assistance in fly-pushing, behavioral observations, and camaraderie during pilot studies. Natasha B. Gallo, Gabriel Moreno-Hagelsieb and Derek Gray provided helpful feed-back and constructive comments, while Michael Steeleworthy of the Wilfrid Laurier University Library is thanked for his help with data archiving. TAFL received support in the for of Natural Sciences and Engineering Research Council (https://www.nserc-crsng.gc.ca) Discovery grants (RGPIN-2016-06133 and RGPIN-2022-03988).This work was conducted at Wilfrid Laurier University, which exists on the traditional territory of the Neutral, Anishnawbe, and Haudenosaunee peoples.

## Notes

***Competing interests*:** The authors declare no competing interests.

https://doi.org/10.5683/SP3/3M8EMU

### Competing Interest Statement

The authors have declared no competing interest.

